# GPR84-Mediated Signal Transduction Promotes Brown Adipocyte Function

**DOI:** 10.1101/2022.10.26.513863

**Authors:** Xuenan Sun, Yu A. An, Vivian A. Paschoal, Camila O. de Souza, May-yun Wang, Lavanya Vishvanath, Lorena Arango, Ayanna Cobb, Joseph A. Nieto Carrion, Harrison Kidd, Shiuhwei Chen, Wenhong Li, Rana K. Gupta, Da Young Oh

## Abstract

G protein-coupled receptor 84 (GPR84), a medium-chain fatty acid receptor, may be involved in various metabolic conditions but the mechanism remains unclear. We found that GPR84 is highly expressed and functions in brown adipose tissue (BAT). GPR84 knockout mice exhibited increased adiposity and vulnerability to cold exposure after aging, along with an increased BAT lipid content and decreased BAT activation compared to wild-type control mice. *In vitro*, primary brown adipocytes from GPR84 knockout mice showed reduced expression of thermogenic genes and lower O_2_ consumption. The GPR84 agonist 6-OAU reversed these effects and restored brown adipocyte activation. *In vivo* and *in vitro* results showed that BAT defects in GPR84 knockout mice were attributed to mitochondrial dysfunction. GPR84 activation greatly affected intracellular calcium efflux, further influencing mitochondrial respiration. GPR84 activates BAT by controlling the mitochondrial calcium content and respiration, suggesting a therapeutic target for activating BAT and treating metabolic diseases.

**HIGHLIGHTS:**

- High expression of GPR84 in brown adipocytes is induced by cold stimulation.
- Aged GPR84 knockout mice exhibit brown adipose tissue defects under cold exposure.
- GPR84 knockout mice show reduced mitochondrial function of brown adipocytes.
- GPR84 activation improves brown adipocyte functions.

## INTRODUCTION

Various adipose depots have gained attention for their roles in regulating systematic energy homeostasis (Kusminski et al., 2016; Mokdad et al., 2003). Among adipose tissues, brown adipose tissue (BAT) can increase energy expenditure by dissipating chemical energy (Cohen and Spiegelman, 2015). Dysfunctions in BAT lead to impaired body temperature homeostasis and glucose handling (Wang et al., 2020). Thus, understanding the pathology of conditions affecting adipose tissue, particularly BAT, may lead to development of effective therapies for metabolic disorders.

Boosting BAT activation is a promising strategy against storing excess energy (Leitner et al., 2017; Rosen and Spiegelman, 2014; van Marken Lichtenbelt et al., 2009). BAT is a thermogenic organ that helps maintain the core body temperature in mammals exposed to cold environments (Porter et al., 2016). In humans, BAT is composed of stromal tissue, white adipose tissue (WAT), and uncoupling protein-1 (UCP1)-containing thermogenic adipocytes (Wu et al., 2012). Upon activation, BAT takes up fatty acids (FAs) and glucose to generate heat (Cypess and Kahn, 2010), thus maintaining the core body temperature. In addition to BAT, beige/brite thermogenic adipocytes are recruited into WAT to respond to stimuli such as cold exposure and β adrenergic receptor-agonist treatment (Lee et al., 2019; Schilperoort et al., 2018). BAT activation after cold exposure also improves whole-body glucose homeostasis and insulin sensitivity in humans and mice (Chondronikola et al., 2014; Poher et al., 2015). Thermogenesis in brown adipocytes mainly involves mitochondria. In fact, mitochondrial dysfunction in BAT leads to metabolic diseases, including insulin resistance, dyslipidemia, and impaired thermogenesis (Johannsen and Ravussin, 2009; Lee *et al*., 2019; Pagliarini and Rutter, 2013). Thus, BAT activation is a promising target for treating metabolic disorders.

G protein-coupled receptors (GPCRs) play critical roles in the regulation of physiological and pathological processes and represent a class of important drug targets. Free FAs act as ligands of multiple GPCRs to activate various signaling pathways (Husted et al., 2017). For example, short-chain FAs (< C6) are recognized by GPR41 and GPR43, which inhibits cAMP production through G_i/o_ signaling (Le Poul et al., 2003; McNelis et al., 2015). Long-chain FAs (> C12) bind to GPR40 to increase intracellular Ca^2+^ levels through G_q/11_ signal transduction, which leads to the activation of insulin secretion in pancreatic beta cells (Fujiwara et al., 2005; Lin et al., 2011). GPR120 is activated by long-chain polyunsaturated FAs, which results in both G_q/11_ and β-arrestin 2-mediated activation of extracellular signal-regulated kinase (ERK) and phosphoatidylinositol-3-kinase (PI3K)/AKT signaling, improves insulin sensitivity, and inhibits inflammation (Ahn et al., 2016; Oh et al., 2010; Wellhauser and Belsham, 2014; Williams-Bey et al., 2014). GPR84 is a medium-chain FA receptor whose function is largely unknown, except macrophages (Hara et al., 2014; Wang et al., 2006). GPR84 is highly expressed in immune cells and is involved in regulating the inflammatory response in macrophages (Du Toit et al., 2018; Recio et al., 2018; Suzuki et al., 2013; Wang *et al*., 2006). Furthermore, GPR84 KO and GPR84 antagonists inhibit Alzheimer’s disease (Audoy-Remus et al., 2015) and diabetic kidney disease by suppressing inflammation (Gagnon et al., 2018). Medium-chain FA levels dynamically change under various metabolic disturbances such as starvation (Steinhauser et al., 2018), feeding (Papamandjaris et al., 1998), cold exposure, type I diabetes (Page et al., 2009), and obesity, indicating that GPR84 activity also varies under these conditions (Page *et al*., 2009; Papamandjaris *et al*., 1998; Schonfeld and Wojtczak, 2016; Steinhauser *et al*., 2018). However, the specific role of GPR84 in these metabolic conditions is unclear. Here, we examined the roles of GPR84 in BAT activation during cold exposure and its effects on mitochondrial function and regulation in brown adipocytes.

## RESULTS

### GPR84 is highly expressed in BAT and brown adipocytes

First, we examined the expression of GPR41, GPR43, GPR120, and GPR84 in human tissues. The expression of these proteins was higher in adipose tissue than in other metabolic tissues, such as pancreas, liver, and muscle (Figures 1A, 1B). Although the functions of GPR43 (FFAR2) (Ge et al., 2008), GPR120 (FFAR4) (Paschoal et al., 2020), and GPR41 (FFAR3) (Amisten et al., 2015) in adipocytes are well-characterized, the role of GPR84 remains unclear. Therefore, we evaluated GPR84 mRNA and protein expression levels in different organs (Figure S1A) and different adipose tissue depots in mice, such as inguinal white adipose tissue (iWAT), gonadal WAT (gWAT), and BAT (Figures 1C, 1D, and Figure S1A). GPR84 was abundantly expressed in BAT (Figures 1C and 1D), suggesting that it plays an important role in this tissue. Because the primary function of BAT is thermoregulation, we examined the expression of GPR84 in the BAT of mice housed at room temperature (23°C) and a cold temperature (6°C) for 7 days. Interestingly, cold exposure induced expression of the GPR84 gene, which was similar to the results observed for UCP1 (Kazak et al., 2017), a marker of BAT activation (Figures 1E and 1F). Furthermore, we isolated the stromal vascular fraction (SVF) of the BAT to induce brown adipocyte differentiation *in vitro*; the expression of GPR84 and UCP1 was gradually increased during differentiation (Figure 1G). Additionally, the expression of GPR84 and UCP1 showed similar patterns in human adipose tissue (Figure S1B). These results indicated that GPR84 is enriched in the BAT and regulates brown adipocyte function.

**Figure 1.**
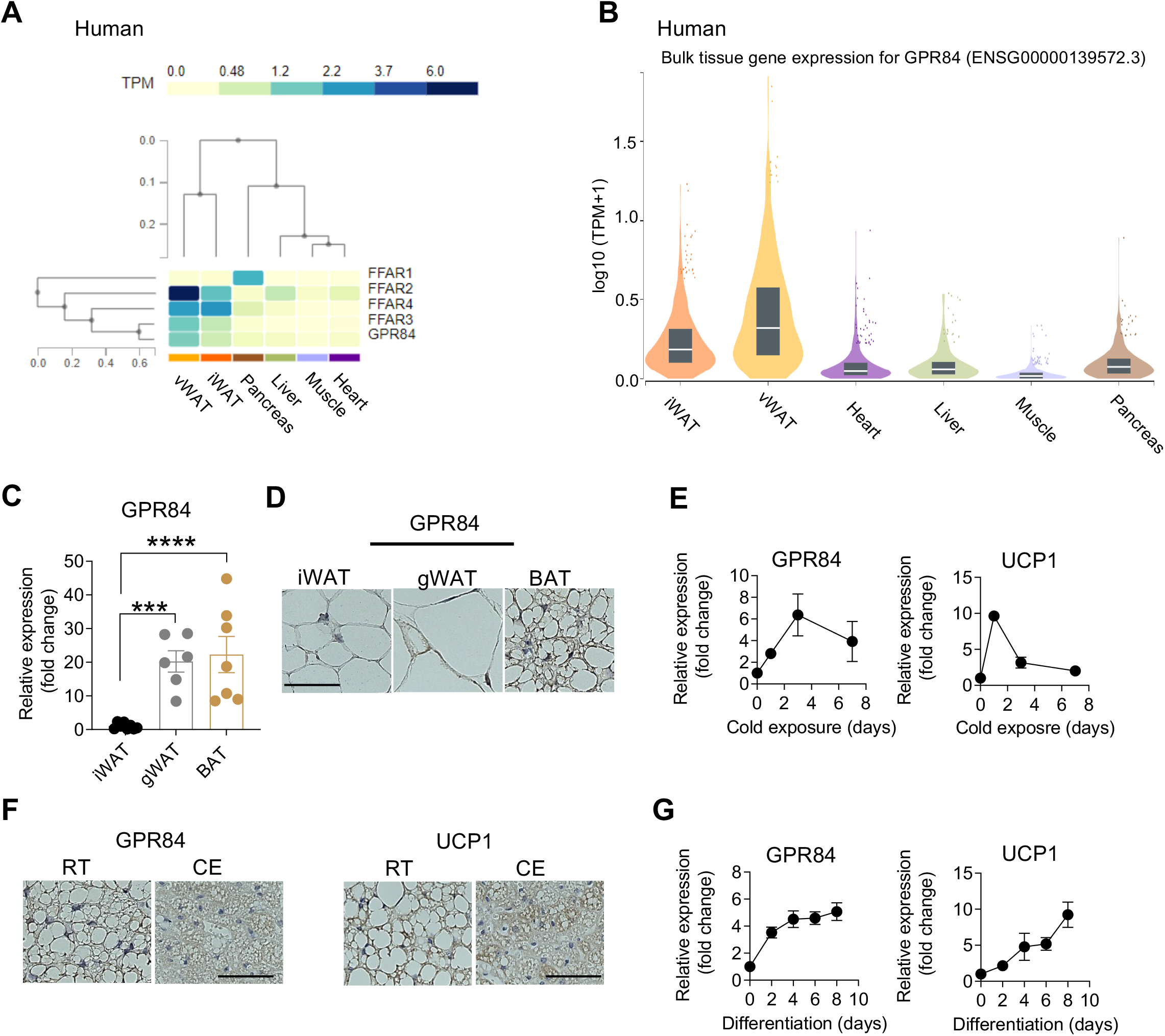
GPR84 is highly expressed in brown adipocytes. **(A)** Expression of different fatty acid receptors in metabolic tissues (vWAT; visceral white adipose tissue, iWAT; inguinal white adipose tissue, pancreas, liver, muscle, and heart) in human samples from the GTEx database (https://gtexportal.org/home/). **(B)** GPR84 expression in different metabolic tissues in human samples from the GTEx database. **(C)** qPCR analysis of GPR84 mRNA expression levels in different adipose tissues, iWAT, gWAT (gonadal WAT), BAT (brown adipose tissue). n=6/group. ****P<0.0001. ***, P < 0.001. **, P < 0.01. *, P < 0.05. Two-tailed Student’s t test was conducted. Data are represented as mean±SEM. **(D)** Representative images (from more than 10 slides/3 different cohorts) of GPR84 immunochemistry staining of different adipose tissues from WT mice. Scale Bar indicates 50 μm. **(E)** qPCR analysis of GPR84 and UCP1 mRNA expression in BAT from WT mice during cold exposure. Triplicate for each time point. n=8-11. **(F)** Representative images (from more than 10 slides/3 different cohorts) of GPR84 and UCP1 staining in BAT from WT mice exposed to cold and room temperature. Scale Bar indicates 50 μm. RT indicates room temperature, and CE indicates cold exposure. **(G)** qPCR analysis of GPR84 and UCP1 mRNA expression during brown adipocyte differentiation from WT mice. Triplicate for each time point. n=8-11. See also Figure S1 and Table S1.

Previous studies indicated that GPR84 activation is involved in the lipopolysaccharide (LPS)-induced inflammatory response (Recio *et al*., 2018). High caloric intake leads to an increase in the incidence of obesity and type 2 diabetes, which is correlated with inflammation (Reilly and Saltiel, 2017). Since the expression of GPR84 was upregulated in the bone marrow and immune cells (Figure S1A, left hand side; Recio *et al*., 2018), we expected the pro-inflammatory role of GPR84 may play a role in the regulation of metabolic phenotype in high fat diet (HFD)-induced obese condition. Therefore, we fed HFD to wild-type (WT) and GPR84 knockout (KO) mice for 16 weeks, and a glucose tolerance test (GTT) and an insulin tolerance test (ITT) were performed. Unexpectedly, glucose homeostasis, insulin sensitivity, and body weight did not differ between the two genotypes on normal chow-fed lean/healthy and HFD-fed obese condition (Figures S1C and S1D). As these findings suggest that GPR84 plays a role in more than just immune cells, we investigated the function of GPR84 in other metabolic tissues, particularly in BAT.

### GPR84 KO leads to BAT defects during aging and cold exposure

To assess the function of GPR84, we compared the phenotype of GPR84 KO mice and WT littermate control mice. Interestingly, aged (13 months old) GPR84 KO mice showed a marked increase in body weight (~10 g) compared to age-matched WT mice (Figures 2A and S2A), although food intake did not differ between the two groups. Body composition analysis revealed that GPR84 KO mice stored more fat mass compared to age-matched WT mice during aging (Figure S2B and S2C). Immunohistochemistry analysis showed a profound unilocular morphology (WAT-like phenotype and decreased expression of UCP1) in the BAT of aged GPR84 KO mice compared to in aged WT mice (Figure 2B). We next analyzed the expression of thermogenic genes in BAT from WT and KO mice at 3 months (young) and 13 months (aged) of age. Quantitative PCR (qPCR) analysis demonstrated that the expression of *UCP1, CideA, PGC1α, Dio2*, and *Cox8b* was downregulated in aged KO mice compared to in aged WT mice, but not in young KO mice (Figure 2C). Analysis of mitochondrial respiration in BAT, iWAT, and gWAT of young (Figure 2D) and aged (Figure 2E) WT and KO mice showed that the oxygen consumption rate (OCR) was ~50% lower in BAT from aged GPR84 KO mice than in BAT from aged WT mice (Figure 2E). However, mitochondrial respiration in WAT did not differ between the two genotypes in young or old mice (Figures S2D–S2G).

**Figure 2.**
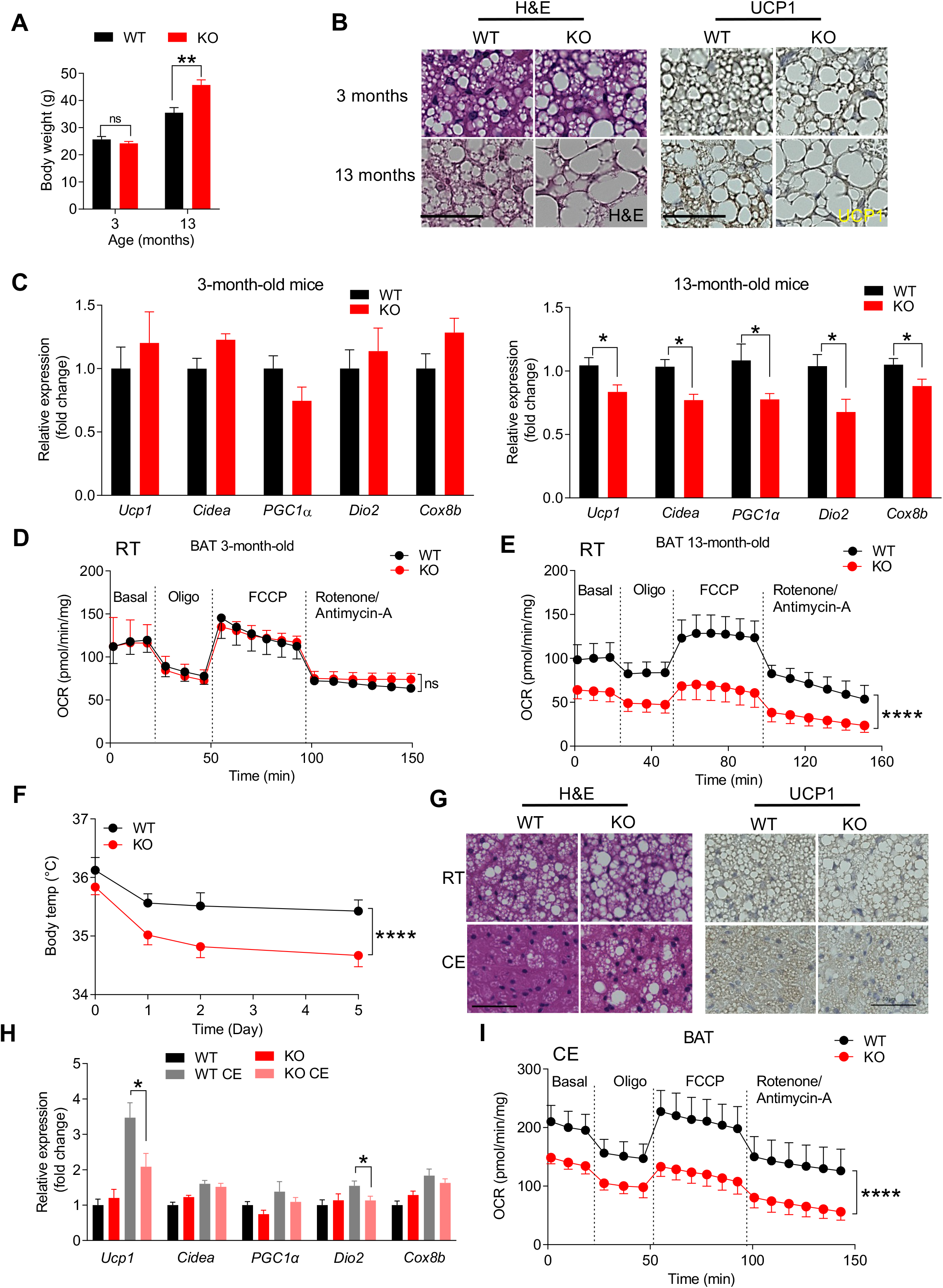
Mice lacking GPR84 leads to BAT dysfunction during aging and cold exposure. **(A)** Body weights of WT and GPR84 KO mice at different ages at RT, n=10/group. **(B)** Representative images of H&E and UCP1 staining in BAT from WT and GPR84 KO mice at different ages. n=10/group. Scale bar indicates 50 μm. **(C)** qPCR analysis of thermogenic gene expressions in BAT from WT and GPR84 KO mice at different ages, n=10/group. **(D)** O_2_ Consumption Rate (OCR) of brown adipose tissue from 3-month-old WT and GPR84 KO mice, n=4. **(E)** O_2_ Consumption Rate (OCR) of brown adipose tissue from 13-month-old WT and GPR84 KO mice, n=4. **(F)** Body temperature of WT and GPR84 KO mice exposed to cold at 3 months of age. **(G)** Representative images of hematoxylin and eosin (H&E) staining and UCP1 immunohistochemistry staining of BAT from WT and GPR84 KO mice exposed to cold at 3 months of age. n=10/group. Scale bar indicates 50 μm. **(H)** qPCR analysis of thermogenic gene expression levels in BAT of WT and GPR84 KO mice, n=10/group. **(I)** O_2_ consumption rate of brown adipose tissue from WT and KO mice after 6 days of cold stimulation, n=3. ****P<0.0001. ***, P < 0.001. **, P < 0.01. *, P < 0.05. CE, cold exposure; RT, room temperature. P values determined by two-tailed Student’s t test; two-way ANOVA followed by a Tukey’s multiple comparison test. Data are represented as mean±SEM. See also Figure S2 and Table S1.

To further evaluate the function of GPR84 in BAT before diverging their body weight between WT and GPR84 KO mice (Figure S2A), we exposed 3 months old, young WT and GPR84 KO mice to cold (6°C). No differences in body weight were found between the groups after 6 days of cold exposure (Figure S2H). Consistently, the weights of different metabolic organs, such as gWAT, BAT, and the liver, had no alterations (Figure S2I). However, the body temperature of GPR84 KO mice was significantly lower than that of WT mice (Figure 2F). Histological analysis revealed that the BAT from KO mice was unilocular (Figure 2G). Thus, the expression of *Ucp1* and *Dio2* was significantly lower in KO mice than in WT mice (Figures 2G and 2H). Furthermore, the expression of WAT-selective genes was higher in BAT from KO mice than in BAT from WT mice after cold exposure (Figure S3A). We then performed a Seahorse experiment to evaluate mitochondrial function and found that OCR in BAT was ~ 50% lower in KO mice than in WT mice after cold adaptation; the OCR showed no differences in gWAT (Figure S3B), iWAT (Figure S3C), and muscle (Figure S3D) between WT and KO mice, indicating that BAT activation was lower in KO mice than in WT mice (Figure 2I). Furthermore, qPCR analysis revealed that the expression of the mitochondrial genes *Apt6*, *Apt8*, and *Nd1* was lower in KO mice than in WT mice (Figures S3E and S3F). Staining of Tim23, a component of the mitochondrial inner membrane import protein, did not differ between the BATs from WT and KO mice after cold exposure, indicating that GPR84 did not affect the numbers of mitochondria following cold exposure (Figure S3G). Serum triglyceride and non-esterified FA (NEFA) levels were dramatically reduced in the BAT of both WT and KO mice after cold exposure. NEFA levels were increased in KO mice after cold exposure compared to the levels in WT mice, suggesting that FA metabolism is dysfunctional in the BAT of KO mice (Figure S3H). Taken together, these findings demonstrate that GPR84 deficiency affects mitochondrial respiration in brown adipocytes, leading to the accumulation of adipose tissue in aged KO mice and defective activation of brown fat in young mice after cold exposure.

### GPR84 activation regulates mitochondrial calcium through the G_i/o_-mediated pathway in brown adipocytes

Next, cultured primary brown adipocytes were used to evaluate whether GPR84 stimulation activates BAT *in vitro*. The immunofluorescence results showed that UCP1 expression was reduced following GPR84 deletion (Figure S4A). Additionally, thermogenic gene expression was significantly downregulated in GPR84 KO brown adipocytes (Figure 3A). Treatment with the GPR84 agonist, 6-n-octylaminouracil (6-OAU) increased lipid accumulation (Figure 3B and 3C) and thermogenic gene expression (Figure 3D) in fully differentiated WT primary brown adipocytes but not in KO brown adipocytes. 6-OAU treatment also increased mitochondrial respiration by ~50% in WT brown adipocytes but did not change the OCR of KO brown adipocytes (Figure 3E). To explore the molecular mechanism by which GPR84 affects brown adipocyte activation, we performed a luciferase reporter activity assay driven by cyclic AMP (cAMP)-responsive element (CRE-luc). As GPR84 is a G_i/o_-coupled receptor that inhibits cAMP (Suzuki *et al*., 2013), HEK 293 cells transfected with GPR84 were first treated with forskolin to increase cAMP levels (Alasbahi and Melzig, 2012) and then treated with different concentrations of 6-OAU. The luciferase assay showed that 6-OAU dose-dependently inhibited the activity of CRE-luc induced by forskolin (Figure S4B). Pertussis toxin (PTX) can inhibit G_i/o_ -dependent signaling pathways; we found that the effect of 6-OAU treatment on CRE-luc activity was abolished in the presence of PTX (Figure S4C). These data indicate that GPR84 primarily activates a PTX-sensitive G_i/o_ pathway to initiate downstream signaling. Activated G_i/o_ proteins release their Gβγ subunits, promoting calcium release from the endoplasmic reticulum (Cheng et al., 2010). The G_i/o_-dependent pathway can affect calcium responses; we found that the luciferase reporter activity driven by serum-responsive element (SRE-luc) in HEK 293 cells transfected with GPR84 was increased by 6-OAU (Figure S4D). Additionally, calcium uptake and efflux through mitochondrial transporters and exchangers can affect mitochondrial function (Giorgi et al., 2018); therefore, we conducted a calcium mobility assay to explore whether GPR84-mediated intracellular calcium modulation affects mitochondrial respiration. We treated WT and KO brown adipocytes with 6-OAU and found that 6-OAU induced a spike of calcium release in WT cells, but not in KO cells (Figure 3F). In the presence of BAPTA-AM, an intracellular calcium chelator, 6-OAU had no effects on WT cells (Figure 3G). As expected, BAPTA-AM completely blocked 6-OAU-induced mitochondrial respiration (Figure S4E). Similarly, the mitochondrial-specific calcium uptake blocker Ru360 (Garcia-Rivas Gde et al., 2006) dramatically inhibited the increase in mitochondrial respiration induced by 6-OAU, indicating that GPR84 activation promotes calcium entry into the mitochondria to increase oxidative phosphorylation. Together, these data demonstrate that GPR84 activation promotes mitochondrial respiration in brown adipocytes via a GPR84 stimulation-mediated increase in the calcium concentration (Figure 3I).

**Figure 3.**
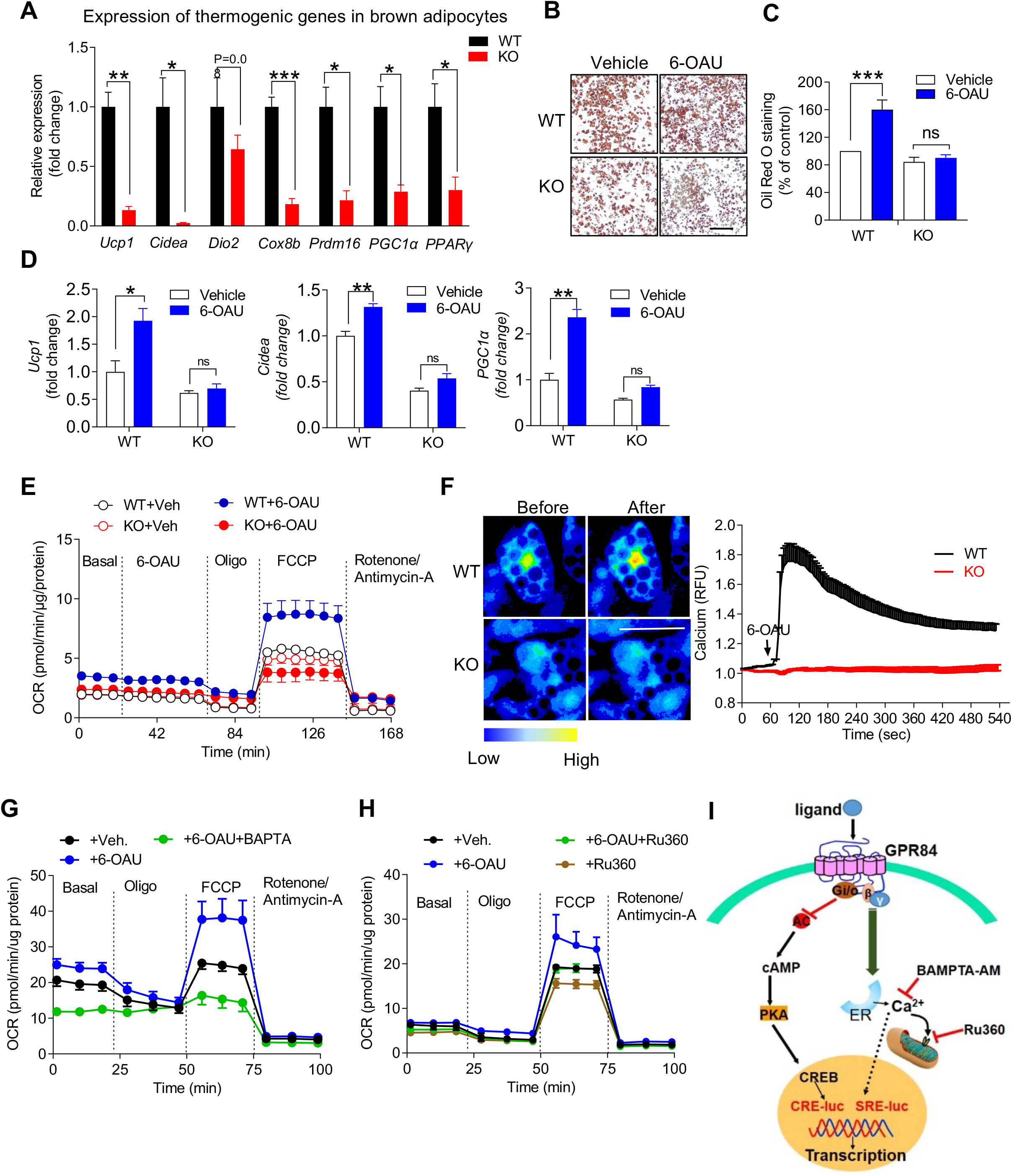
Activated GPR84 regulates mitochondrial calcium through Gi/o-mediated pathway in brown adipocytes. **(A)** qPCR analysis of thermogenic gene expression in SVF differentiated brown adipocytes from WT and GPR84 KO mice. **(B and C) (B)** Representative images of lipid staining in WT and KO differentiated brown adipocytes with or without 6-OAU. **(C)** quantification of (B). Scale Bar indicates 50 μm. **(D)** qPCR analysis of thermogenic gene levels in WT and KO brown adipocytes ± 6-OAU. **(E)** Mitochondrial respiration at different time points in WT and KO brown adipocytes. Cells were pretreated ± 6-OAU for 30 min, before oxygen consumption rate (OCR) was measured in a seahorse X24 analyzer. **(F)** WT and GPR84 KO brown adipocytes were incubated with the calcium-sensitive dye Fluo-4-AM for 1 h at RT, followed by live cell imaging with a confocal laser scanning microscope (LSM 510, Zeiss) and stimulation with 6-OAU (50 μM). Scale bar indicates 50μm. Relative fluorescence units were analyzed by ROI in Image J. **(G)** Brown adipocytes were pretreated with 6-OAU (50 μM) for 1h, and then treated with BAPTA-AM for 30 minutes, after which the OCR was determined in a seahorse X24 analyzer. n=5/each time point. **(H)** Brown adipocytes were pretreated with 6-OAU (50 μM) for 1h, and then treated with Ru360 for another 1h, after which the OCR was determined in a seahorse X24 analyzer. n=5/each time point. **(I)** Schematic diagram for the molecular mechanism of the GPR84-mediated mitochondrial respiration, left letters and arrows indicate the G_i_-mediated CRE-luc inhibition, and right colored letters and arrows indicate the Gβγ-induced calcium pathway *P < 0.05, **P < 0.01. P values determined by two-tailed Student’s t test; two-way ANOVA followed by a Tukey’s multiple comparison test. Data are represented as mean±SEM. See also Figure S3 and Table S1.

### GPR84 agonist 6-OAU promotes BAT activation in mice exposed to cold

To evaluate the effect of GPR84 activation *in vivo*, the mice were infused with 6-OAU using an osmotic minipump and then subjected to cold stress (Figure 4A). The body and metabolic tissue weights were not altered after 6 days of cold exposure (Figure 4B). However, mice infused with 6-OAU maintained their core body temperature better compared to mice infused with vehicle (Figure 4B). The immunohistochemistry data revealed more multilocular brown adipocytes in the BAT of 6-OAU-infused mice than in that of vehicle-infused control mice (Figure 4C). UCP1 staining showed that cold-induced BAT activation was stronger in 6-OAU-treated mice than in control mice (Figure 4D). Consistently, expression of thermogenic genes, including *Ucp1*, *PGC1α*, and *PPARγ*, was upregulated in the BAT of 6-OAU-infused mice (Figure 4E). Furthermore, the OCR in the BAT of mice infused with 6-OAU was ~50% higher than that in the BAT of control mice (Figure 4F). However, there were no changes in the OCR of iWAT (Figure 4G) and gWAT (Figure 4H) in any of the groups. Short-term injection of 6-OAU can increase inflammation in rats (Suzuki *et al*., 2013); therefore, we examined whether the levels of inflammatory cytokines such as TNFα, IL-1β, IL-6, and MCP1 changed after 6-OAU infusion. No differences were found in the levels of the inflammatory cytokines (Figure 4I), indicating that the effect of 6-OAU on BAT activation was independent of inflammation. Similarly, the expression of these inflammatory genes in BAT did not differ between groups (Figure 4I), suggesting that 6-OAU infusion does not affect inflammation in these mice. Collectively, these findings show that GPR84 activation specifically affects BAT activation and, consequently, the maintenance of body temperature during cold exposure.

**Figure 4.**
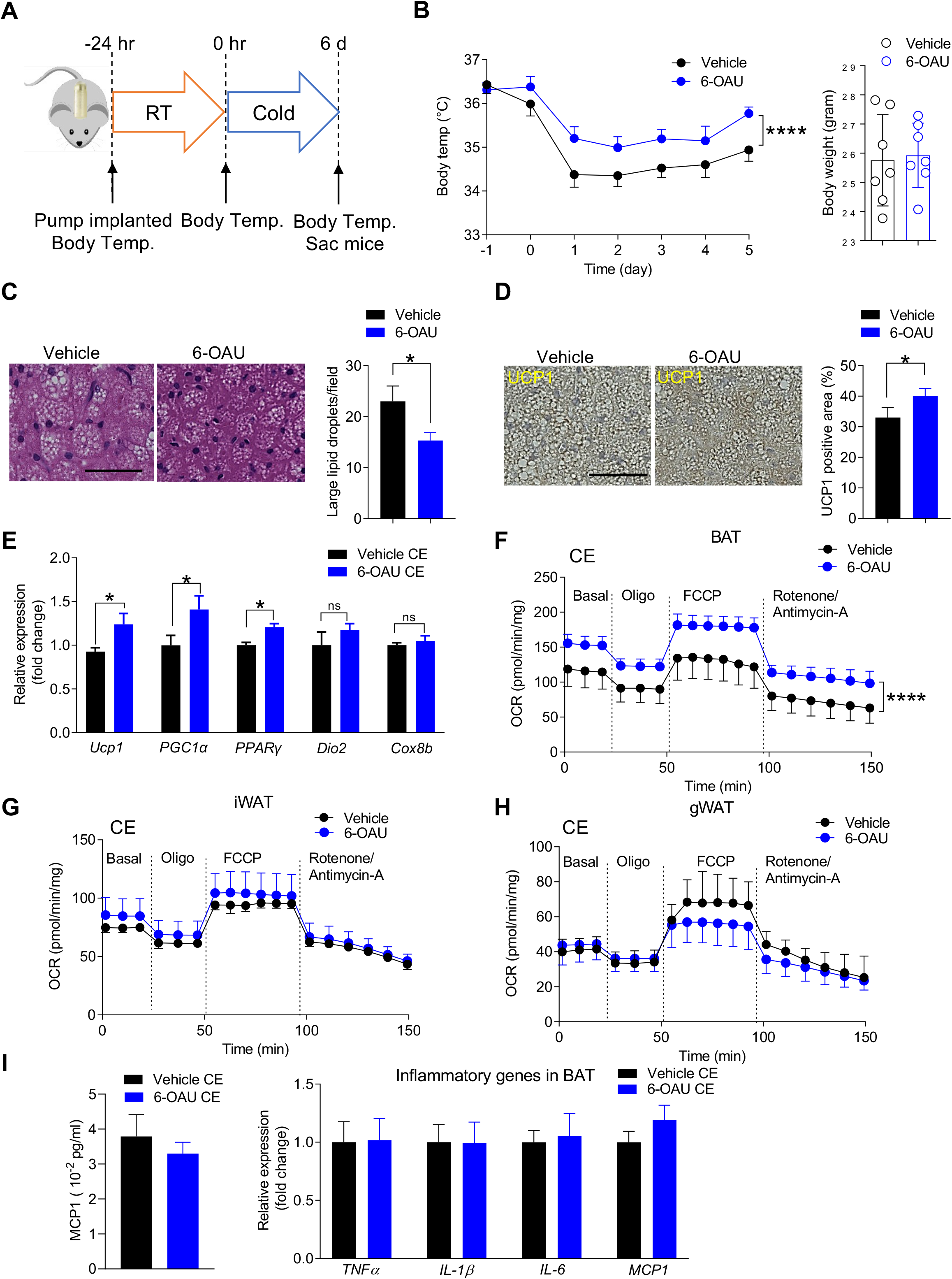
GPR84 agonist 6-OAU promotes BAT activation in mice on cold exposure. **(A)** Experimental design for 6-OAU treatment in mice, n=10 per group. **(B)** Body temperature and body weight of Vehicle and 6-OAU treated mice exposed to cold, n=10/group. **(C)** Representative images of hematoxylin and eosin (H&E) staining of BAT from 6-OAU treated mice, scale bar 50 indicates μm and quantification of lipid droplets for H&E staining, n=3/group. **(D)** Representative UCP1 immunohistochemistry staining in BAT from vehicle and 6-OAU treated mice, n=4/group (**E)** qPCR analysis of mRNA levels of thermogenic genes in BAT of vehicle and 6-OAU treated mice, n=8-10/group. **(F)** O_2_ consumption rate of brown adipose tissue from 6-OAU and vehicle treated mice after 6 days of cold stimulation, n=3. **(G and H)** O_2_ consumption rate for iWAT (G) and gWAT (H) from 6-OAU treated mice on cold exposure. n=4/group. **(I)** plasma level of MCP1 in control and 6-OAU treated mice were measured by ELISA and inflammatory gene expressions in BAT from 6-OAU infused mice after cold stimulation. n=8-10/group. *P < 0.05, **P < 0.01. CE, cold exposure; RT, room temperature. P values determined by two-tailed Student’s t test; two-way ANOVA followed by a Tukey’s multiple comparison test. Data are represented as mean±SEM. See also Figure S4, Table S1 and Table S2.

## DISCUSSION

Targeting signaling pathways in adipose tissue represents a promising approach to controlling their functions (Fuster et al., 2016; Kusminski *et al*., 2016; Reilly and Saltiel, 2017). Although GPR84 is highly expressed in the skeletal muscle and adipose tissue, its deficiency does not affect the inflammatory status in skeletal muscle (Montgomery et al., 2019), liver (Simard et al., 2020), or adipose tissue, suggesting that GPR84 is not involved in inflammatory regulation in these tissues. The function of GPR84 in adipocytes is unclear. Here, we found that GPR84 is highly expressed in human and mouse adipose tissue, and its expression in BAT increases with cold stimulation and during the maturation of brown adipocytes. These findings led us to explore the role of GPR84 in BAT. We demonstrated that GPR84 plays a crucial role in BAT activation under cold stimulation and aging, whereas GPR84 KO mice showed cold intolerance and increased body weights during aging. Furthermore, GPR84 activation promoted brown adipocyte activity through G_i/o_-dependent calcium activation and improved BAT function after cold exposure. Collectively, our data indicate that GPR84 is a promising target for BAT stimulation to improve metabolic disorders.

Aging-related dysfunction in adipose tissues may lead to age-related metabolic alterations (Lopez-Otin et al., 2013). Aging is characterized by an increase in adiposity and a decrease in BAT depots and activity as well as UCP1 expression. Various factors may influence age-associated involution of BAT, including the loss of mitochondrial function, impairment of the sympathetic nervous system, age-induced alteration of brown adipogenic stem/progenitor cell function, and changes in endocrine signals (Zoico et al., 2019). In this study, we demonstrated that GPR84 KO mice accelerated aging-mediated body weight gain and BAT dysfunction compared to WT mice. Aged GPR84 KO mice showed impaired mitochondrial function (impaired OCR) and reduced thermogenic genes in BAT compared to the BAT of WT mice, suggesting that GPR84 activation during aging can attenuate the functional decline of the BAT. We determined the specific age (3-month-old) at which the body weight had not been diverged yet but showed clear differences in the BAT phenotype in WT and KO mice to assess the role of GPR84 in BAT. Mitochondrial dysfunction is involved in the pathogenesis of several age-related disorders, such as type 2 diabetes and obesity (Poher *et al*., 2015; Sastre et al., 2003); thus, mitochondrial dysfunction in young GPR84 KO mice may have been a driving cause and exerted accumulating effects that accelerated aging phenotypes in GPR84 KO mice compared to in WT control mice.

Adaptive thermogenesis has attracted attention because of its ability to increase systemic energy expenditure and counter obesity and diabetes (Carobbio et al., 2019; Harms and Seale, 2013; Rosen and Spiegelman, 2014). In thermogenesis, BAT is essential for classical non-shivering thermogenesis as well as for cold acclimation-recruited norepinephrine-induced thermogenesis. β_3_-Adrenergic signaling is the dominant signaling pathway controlling non-shivering thermogenesis in brown and beige adipocytes (Ahmad et al., 2021). Upon cold exposure, norepinephrine released from the sympathetic nervous system binds mainly to the β_3_-adrenergic receptor to induce adrenergic signaling (Liu et al., 2019; Schena and Caplan, 2019). The β_3_-adrenergic receptor coupled with G_s_ activates adenylate cyclase, which in turn produces cAMP for protein kinase A activation. Upon β_3_-adrenergic receptor activation, protein kinase A phosphorylates hormone-sensitive lipase as well as perilipin-1 on lipid droplets to promote lipolysis. Thus, FAs produced upon lipolysis can undergo β-oxidation to eventually produce NADH and FADH for use in the electron transport chain during UCP1-mediated thermogenesis (Shah et al., 2013). FAs also directly bind to UCP1 to facilitate proton influx into the mitochondrial matrix (Kajimura and Saito, 2014). In contrast, in a fed state, FAs as substrates for thermogenesis appear to originate from lipolysis of WAT, although it was also reported that inhibition of intracellular lipolysis suppressed cold-induced non-shivering thermogenesis in BAT, which was compensated by an increased shivering in humans (Blondin et al., 2017; Shin et al., 2017). New thermogenesis pathways in adipose tissues have been discovered in the past few years. Ikeda et al. identified a non-canonical thermogenic mechanism through which beige fat controls whole-body energy homeostasis via Ca^2+^ cycling (Ikeda et al., 2017; Long et al., 2016). GPR84 signaling through a G_i/o_-coupled pathway activates calcium release from the endoplasmic reticulum (Cheng *et al*., 2010). Our results revealed that GPR84 activation induces intracellular calcium release, which promotes mitochondrial respiration in primary brown adipocytes. The mitochondrial morphology, depolarization, fission, and fusion are coupled to calcium mobilization (Barsukova et al., 2011; Iglewski et al., 2010); however, the precise mechanism of the effects of GPR84 stimulation on mitochondrial dynamics and structure requires further analysis.

Our results of 6-OAU-infused mice are quite interesting and consistent with those of the KO phenotype. GPR84 activation *in vivo* improved BAT function, while 6-OAU infusion did not increase Inflammation (Figure 4I). The concentration we used for 6-OAU infusion activated GPR84 in the BAT without the involvement of inflammatory factors such as CXCL-1; a previous study showed that high-dose of short-term 6-OAU injection increased CXCL1 levels (Suzuki *et al*., 2013). Thus, GPR84 activation is not involved in inflammatory regulation in the BAT.

Taken together, our findings demonstrate that GPR84 deficiency accelerated aging-mediated BAT defects and body weight gain, and that GPR84 activation-mediated calcium signaling stimulates the function of BAT in controlling thermogenesis.

## LIMITATIONS OF THE STUDY

We demonstrated that GPR84 activation by 6-OAU promotes mitochondrial respiration in brown adipocytes through G_i/o_-mediated increase of intracellular calcium concentration. 6-OAU infusion *in vivo* successfully activated brown adipose tissue and thermogenesis as we observed *in vitro* experiments. Although GPR84 is known as a receptor for medium-chain FAs, we only used 6-OAU as a selective agonist for GPR84 throughout this study due to the lower sensitivity of medium-chain FAs compared to 6-OAU. Since medium-chain FAs are generated through the metabolic regulation and they are endogenous ligands for GPR84, it would be better reflected the physiological role of GPR84 activation by medium-chain FAs *in vivo*. Future studies to utilize medium-chain FAs to activate GPR84 *in vivo* by direct infusion and/or feeding medium-chain FAs containing diet to WT and GPR84 KO mice will be needed.

## AUTHOR CONTRIBUTIONS

X.S. and D.Y.O. conceived and designed the study. X.S. performed most of the experiments with helps from Y.A.A. and M.W. (Seahorse experiments); V.P., C.S., A.C., J.A.N.C., S.C., and L.A. (mouse surgeries and tissue processing for RNA isolation and immunohistochemistry); H.K. and W.L. (calcium mobility assay); and L.V. and R.K.G. (primary adipocyte culture). R.K.G. provided the key reagent and discussed results. X.S. and D.Y.O. analyzed, interpreted data, and co-wrote this manuscript.

## ACKNOWLEDGEMENTS

We thank the UTSW Molecular Pathology Core (John Shelton) for assistance with histology analysis. We also thank the Animal Resources Center of UTSW for mouse generation, breeding, and care. This work was supported by grants from the NIH NIDDK (R01 DK108773 to D.Y.O.; 5R01 DK104789 to R.K.G.).

## DECLARATION OF INTERESTS

The authors declare no competing financial interests.

## STAR METHODS

### KEY RESOURCES TABLE

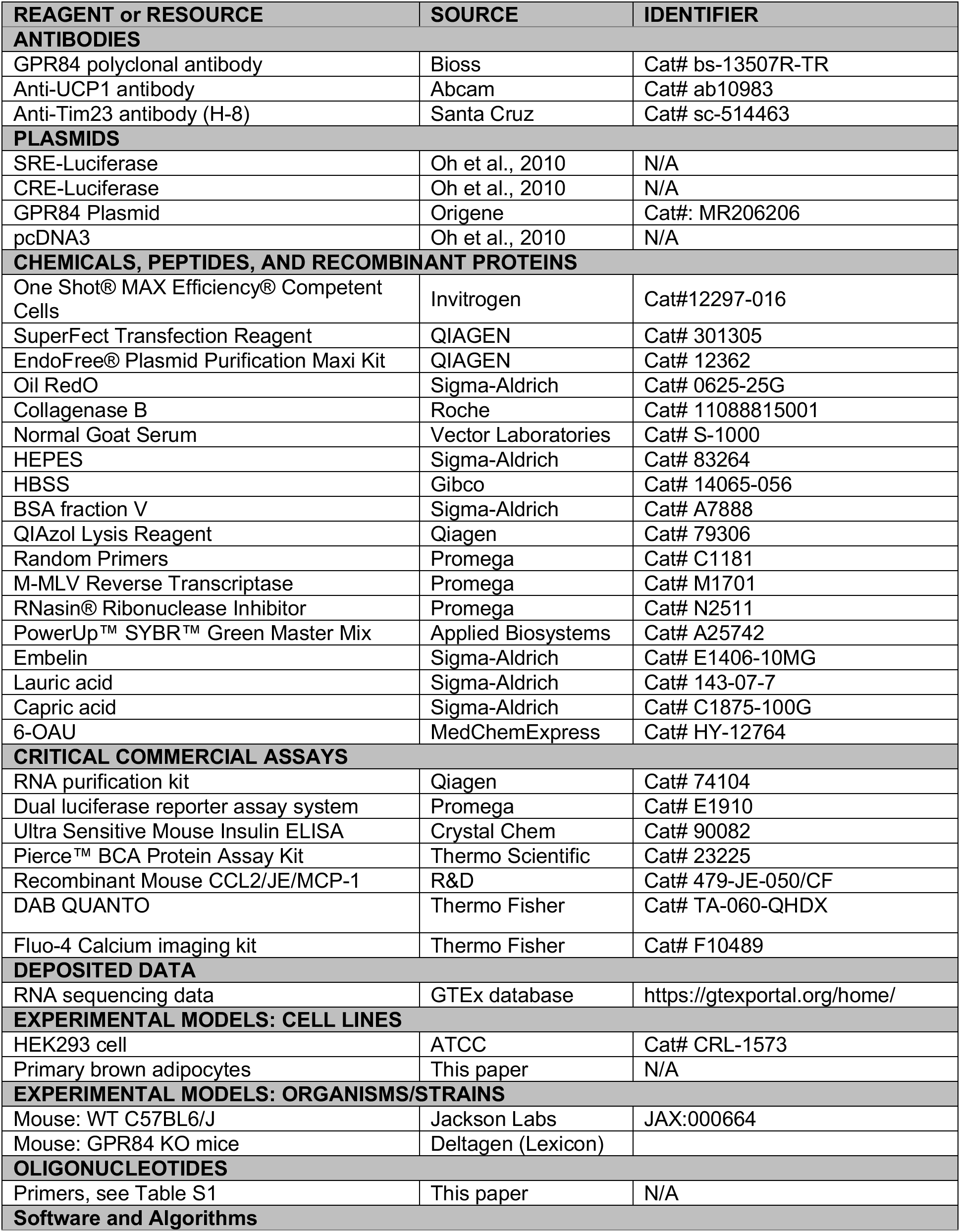
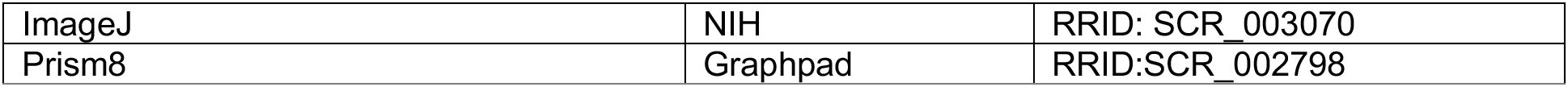

### LEAD AND MATERIALS AVAILABILITY

Further information and requests for resources and reagents should be directed and will be fulfilled by the corresponding author, Da Young Oh, PhD (UT Southwestern Medical Center).

### EXPERIMENTAL MODEL AND SUBJECT DETAILS

#### Animal care and use

GPR84 knockout (KO) mice were bought from Deltagene and housed at specific pathogen-free facility (UT Southwestern Medical Center). Male C57Bl/6 (wild type, WT) or GPR84 KO littermates, from 8 weeks of age, were fed a normal chow (13.5% fat; LabDiet) or high-fat diet (60% fat; Cat# D12492; Research Diet) provided *ad libitum* for 14 weeks. Mice received fresh diet weekly, and food consumption and body weight were monitored. 12-week-old mice were housed at 6°C for 6 days. During cold exposure, mouse body temperature was monitored every one or two days.

#### Study approval

All procedures on animals have been approved by the Institutional Animal Care and Use Committee (IACUC) of UT Southwestern Medical Center (APN no. 2016-101841-G).

### METHOD DETAILS

#### Plasma triglycerides measurement

After mouse euthanization, analysis of plasma total cholesterol, triglycerides and free-fatty-acid (NEFA) was performed on whole blood samples obtained by venipuncture in tubes and centrifuged at 3,000 r.c.f. for 15 min. Samples were analyzed through a Vitros 250 Chemistry system (Ortho Clinical Diagnostics).

#### Calcium mobility assay

Differentiated brown adipocytes were incubated with calcium sensitive dye Fluo-4-AM for 15 min at 37°C and another 15 min at room temperature in DMEM without FBS (Schilperoort *et al*., 2018). After washing cells with live cell image solution twice, cells were ready to perform assay. By using a confocal laser scanning microscope, photos were taken every 4 seconds in the presence or absence of 6-OAU in both WT and GPR84 KO cells. Images and videos were recorded and analyzed by Openlab software.

#### RNA isolation, cDNA synthesis and qPCR

Total RNA was extracted from brown adipose tissue, primary brown adipocytes using an RNA purification kit (QIAGEN). The cDNA was synthesized using SuperScript III and random primers. Quantitative PCR was carried out in 10 μL reactions with SYBR Green master mix (Applied Biosystems) on ABI Real-Time PCR System (Applied Biosystems). Relative gene expression was normalized to the standard housekeeping gene (RPL19) using the ΔΔCT method. The specificity of the PCR amplification was verified by melting curve analysis of the final products using ABI software. Primer sequences are available in *Table S1*.

#### Cell culture

Brown pre-adipocytes were cultured in DMEM (Corning) supplemented with 10% fetal bovine serum (FBS), penicillin–streptomycin and gentamicin (no. 15750060, Gibco, ThermoFisher Scientific) in a humid incubator with 10% CO_2_ at 37°C. Pre-adipocytes were differentiated with induction medium including 0.5 mM 3-isobutyl-1-methylxanthine (IBMX), 1 μM dexamethasone, 5 μg/mL insulin to initiate in vitro differentiation for 2 days after the cells reached over 95% confluence. After 48h of incubation, cells were cultured with maintenance medium containing only 5 μg/mL insulin for 6 days.

#### Tissue histology and immunohistochemistry

Mouse brown adipose tissues were paraffin embedded and were cut in 4 μm sections and stained with hematoxylin and eosin (H&E), GPR84 and UCP1, performed by the University of Texas Southwestern Medical Center Histology Core. Images (×100 or ×200 magnification) were acquired by the FSX100 Inverted Microscope (Olympus), and representative histological images are shown. Paraffin-embedded slides were prepared by the University of Texas Southwestern Medical Center Histology Core for immunohistochemistry staining. Briefly, deparaffinized sections were stained with anti-GPR84 (1:50), anti-UCP1 (1:500) primary antibodies and incubated overnight at 4°C. Then the secondary antibodies (Life Technologies) were added for 2 h, and finally the cover slips were added.

#### Cellular oxygen consumption measurements

Mitochondrial respiration was examined by using the Seahorse XF24 Extracellular Flux Analyzer (Agilent) according to the manufacturers’ instructions. Briefly, a mitochondrial respiration was determined following a manufacturer-recommended BOFA (basal-oligomycin-FCCP-antimycin A/rotenone) protocol. Ex vivo and *in vitro* mitochondrial function were measured by utilizing 3-5 mg brown fat tissue and SVF-differentiated brown adipocytes, respectively. For tissues and cultured cells, 6-OAU (50 μM), oligomycin (2 μM), FCCP (8 μM) and antimycin A (10 μM) plus rotenone (3 μM) were injected. OCR was acquired through the Seahorse instrument.

#### Oil red O staining

Differentiated brown adipocytes were fixed with 4% paraformaldehyde (15 min, RT) and rinsed with PBS 3 times. Cells were stained with 0.15% Oil-Red O solution (60 min, RT), after which they were washed with ddH_2_O 4 times. The cells were ready for taking pictures.

#### Body composition analysis

Body weight of mice were measured with a scale, and total body lean and fat mass with the Bruker Minispec mq10 system (Bruker).

#### 6-OAU infusion in mice and body temperature monitor

Minipumps (1007D; Alzet, Cupertino, CA) were implanted subcutaneously in 11- to 12-week-old mice to deliver 6-OAU (6 mg/kg per day) or vehicle (PBS). For body temperature measurements, mice were infused with 6-OAU or vehicle for 2 weeks and Body temperature was measured using Analysis System (Visitech Systems, Napa Place Apex, NC). Minipumps were implanted 7 days later to infuse 6-OAU or vehicle.

### QUANTIFICATION AND STATISTICAL ANALYSIS

Data are presented as the mean ± SEM. The significance of differences between groups was evaluated using analysis of variance. The *P* value < 0.05 was considered significant. Statistical parameters including the exact value of *n*, the definition of center, dispersion and precision measures (mean ± SEM) and statistical significance are reported in the Figures and the Figure Legends. Data is judged to be statistically significant when *P* < 0.05 by two-tailed Student’s t-Test or two-way ANOVA followed by a Tukey’s multiple comparison test, where appropriate. In Figures, asterisks denote statistical significance (*, *P* < 0.05). Statistical analysis was performed in GraphPad PRISM 8.

## DATA AND SOFTWARE AVAILABILITY

### Software availability

The software is available on the web sites linked in the Key Resources Table.

## Notes

### Competing Interest Statement

The authors have declared no competing interest.

## REFERENCES

Ahmad, A., Naqvi, S.A., Jaskani, M.J., Waseem, M., Ali, E., Khan, I.A., Faisal Manzoor, M., Siddeeg, A., and Aadil, R.M. (2021). Efficient utilization of date palm waste for the bioethanol production through Saccharomyces cerevisiae strain. Food Sci Nutr 9, 2066–2074. 10.1002/fsn3.2175.

Ahn, S.H., Park, S.Y., Baek, J.E., Lee, S.Y., Baek, W.Y., Lee, S.Y., Lee, Y.S., Yoo, H.J., Kim, H., Lee, S.H., et al. (2016). Free Fatty Acid Receptor 4 (GPR120) Stimulates Bone Formation and Suppresses Bone Resorption in the Presence of Elevated n-3 Fatty Acid Levels. Endocrinology 157, 2621–2635. 10.1210/en.2015-1855.

Alasbahi, R.H., and Melzig, M.F. (2012). Forskolin and derivatives as tools for studying the role of cAMP. Pharmazie 67, 5–13.

Amisten, S., Neville, M., Hawkes, R., Persaud, S.J., Karpe, F., and Salehi, A. (2015). An atlas of G-protein coupled receptor expression and function in human subcutaneous adipose tissue. Pharmacol Ther 146, 61–93. 10.1016/j.pharmthera.2014.09.007.

Audoy-Remus, J., Bozoyan, L., Dumas, A., Filali, M., Lecours, C., Lacroix, S., Rivest, S., Tremblay, M.E., and Vallieres, L. (2015). GPR84 deficiency reduces microgliosis, but accelerates dendritic degeneration and cognitive decline in a mouse model of Alzheimer’s disease. Brain Behav Immun 46, 112–120. 10.1016/j.bbi.2015.01.010.

Barsukova, A.G., Bourdette, D., and Forte, M. (2011). Mitochondrial calcium and its regulation in neurodegeneration induced by oxidative stress. Eur J Neurosci 34, 437–447. 10.1111/j.1460-9568.2011.07760.x.

Blondin, D.P., Frisch, F., Phoenix, S., Guerin, B., Turcotte, E.E., Haman, F., Richard, D., and Carpentier, A.C. (2017). Inhibition of Intracellular Triglyceride Lipolysis Suppresses Cold-Induced Brown Adipose Tissue Metabolism and Increases Shivering in Humans. Cell Metab 25, 438–447. 10.1016/j.cmet.2016.12.005.

Carobbio, S., Guenantin, A.C., Samuelson, I., Bahri, M., and Vidal-Puig, A. (2019). Brown and beige fat: From molecules to physiology and pathophysiology. Biochim Biophys Acta Mol Cell Biol Lipids 1864, 37–50. 10.1016/j.bbalip.2018.05.013.

Cheng, Z., Garvin, D., Paguio, A., Stecha, P., Wood, K., and Fan, F. (2010). Luciferase Reporter Assay System for Deciphering GPCR Pathways. Curr Chem Genomics 4, 84–91. 10.2174/1875397301004010084.

Chondronikola, M., Volpi, E., Borsheim, E., Porter, C., Annamalai, P., Enerback, S., Lidell, M.E., Saraf, M.K., Labbe, S.M., Hurren, N.M., et al. (2014). Brown adipose tissue improves whole-body glucose homeostasis and insulin sensitivity in humans. Diabetes 63, 4089–4099. 10.2337/db14-0746.

Cohen, P., and Spiegelman, B.M. (2015). Brown and Beige Fat: Molecular Parts of a Thermogenic Machine. Diabetes 64, 2346–2351. 10.2337/db15-0318.

Cypess, A.M., and Kahn, C.R. (2010). The role and importance of brown adipose tissue in energy homeostasis. Curr Opin Pediatr 22, 478–484. 10.1097/MOP.0b013e32833a8d6e.

Du Toit, E., Browne, L., Irving-Rodgers, H., Massa, H.M., Fozzard, N., Jennings, M.P., and Peak, I.R. (2018). Effect of GPR84 deletion on obesity and diabetes development in mice fed long chain or medium chain fatty acid rich diets. Eur J Nutr 57, 1737–1746. 10.1007/s00394-017-1456-5.

Fujiwara, K., Maekawa, F., and Yada, T. (2005). Oleic acid interacts with GPR40 to induce Ca2+ signaling in rat islet beta-cells: mediation by PLC and L-type Ca2+ channel and link to insulin release. Am J Physiol Endocrinol Metab 289, E670–677. 10.1152/ajpendo.00035.2005.

Fuster, J.J., Ouchi, N., Gokce, N., and Walsh, K. (2016). Obesity-Induced Changes in Adipose Tissue Microenvironment and Their Impact on Cardiovascular Disease. Circ Res 118, 1786–1807. 10.1161/CIRCRESAHA.115.306885.

Gagnon, L., Leduc, M., Thibodeau, J.F., Zhang, M.Z., Grouix, B., Sarra-Bournet, F., Gagnon, W., Hince, K., Tremblay, M., Geerts, L., et al. (2018). A Newly Discovered Antifibrotic Pathway Regulated by Two Fatty Acid Receptors: GPR40 and GPR84. Am J Pathol 188, 1132–1148. 10.1016/j.ajpath.2018.01.009.

Garcia-Rivas Gde, J., Carvajal, K., Correa, F., and Zazueta, C. (2006). Ru360, a specific mitochondrial calcium uptake inhibitor, improves cardiac post-ischaemic functional recovery in rats in vivo. Br J Pharmacol 149, 829–837. 10.1038/sj.bjp.0706932.

Ge, H., Li, X., Weiszmann, J., Wang, P., Baribault, H., Chen, J.L., Tian, H., and Li, Y. (2008). Activation of G protein-coupled receptor 43 in adipocytes leads to inhibition of lipolysis and suppression of plasma free fatty acids. Endocrinology 149, 4519–4526. 10.1210/en.2008-0059.

Giorgi, C., Marchi, S., and Pinton, P. (2018). The machineries, regulation and cellular functions of mitochondrial calcium. Nat Rev Mol Cell Biol 19, 713–730. 10.1038/s41580-018-0052-8.

Hara, T., Kashihara, D., Ichimura, A., Kimura, I., Tsujimoto, G., and Hirasawa, A. (2014). Role of free fatty acid receptors in the regulation of energy metabolism. Biochim Biophys Acta 1841, 1292–1300. 10.1016/j.bbalip.2014.06.002.

Harms, M., and Seale, P. (2013). Brown and beige fat: development, function and therapeutic potential. Nat Med 19, 1252–1263. 10.1038/nm.3361.

Husted, A.S., Trauelsen, M., Rudenko, O., Hjorth, S.A., and Schwartz, T.W. (2017). GPCR-Mediated Signaling of Metabolites. Cell Metab 25, 777–796. 10.1016/j.cmet.2017.03.008.

Iglewski, M., Hill, J.A., Lavandero, S., and Rothermel, B.A. (2010). Mitochondrial fission and autophagy in the normal and diseased heart. Curr Hypertens Rep 12, 418–425. 10.1007/s11906-010-0147-x.

Ikeda, K., Kang, Q., Yoneshiro, T., Camporez, J.P., Maki, H., Homma, M., Shinoda, K., Chen, Y., Lu, X., Maretich, P., et al. (2017). UCP1-independent signaling involving SERCA2b-mediated calcium cycling regulates beige fat thermogenesis and systemic glucose homeostasis. Nat Med 23, 1454–1465. 10.1038/nm.4429.

Johannsen, D.L., and Ravussin, E. (2009). The role of mitochondria in health and disease. Curr Opin Pharmacol 9, 780–786. 10.1016/j.coph.2009.09.002.

Kajimura, S., and Saito, M. (2014). A new era in brown adipose tissue biology: molecular control of brown fat development and energy homeostasis. Annu Rev Physiol 76, 225–249. 10.1146/annurev-physiol-021113-170252.

Kazak, L., Chouchani, E.T., Stavrovskaya, I.G., Lu, G.Z., Jedrychowski, M.P., Egan, D.F., Kumari, M., Kong, X., Erickson, B.K., Szpyt, J., et al. (2017). UCP1 deficiency causes brown fat respiratory chain depletion and sensitizes mitochondria to calcium overload-induced dysfunction. Proc Natl Acad Sci U S A 114, 7981–7986. 10.1073/pnas.1705406114.

Kusminski, C.M., Bickel, P.E., and Scherer, P.E. (2016). Targeting adipose tissue in the treatment of obesity-associated diabetes. Nat Rev Drug Discov 15, 639–660. 10.1038/nrd.2016.75.

Le Poul, E., Loison, C., Struyf, S., Springael, J.Y., Lannoy, V., Decobecq, M.E., Brezillon, S., Dupriez, V., Vassart, G., Van Damme, J., et al. (2003). Functional characterization of human receptors for short chain fatty acids and their role in polymorphonuclear cell activation. J Biol Chem 278, 25481–25489. 10.1074/jbc.M301403200.

Lee, J.H., Park, A., Oh, K.J., Lee, S.C., Kim, W.K., and Bae, K.H. (2019). The Role of Adipose Tissue Mitochondria: Regulation of Mitochondrial Function for the Treatment of Metabolic Diseases. Int J Mol Sci 20. 10.3390/ijms20194924.

Leitner, B.P., Huang, S., Brychta, R.J., Duckworth, C.J., Baskin, A.S., McGehee, S., Tal, I., Dieckmann, W., Gupta, G., Kolodny, G.M., et al. (2017). Mapping of human brown adipose tissue in lean and obese young men. Proc Natl Acad Sci U S A 114, 8649–8654. 10.1073/pnas.1705287114.

Lin, D.C., Zhang, J., Zhuang, R., Li, F., Nguyen, K., Chen, M., Tran, T., Lopez, E., Lu, J.Y., Li, X.N., et al. (2011). AMG 837: a novel GPR40/FFA1 agonist that enhances insulin secretion and lowers glucose levels in rodents. PLoS One 6, e27270. 10.1371/journal.pone.0027270.

Liu, J., Wang, Y., and Lin, L. (2019). Small molecules for fat combustion: targeting obesity. Acta Pharm Sin B 9, 220–236. 10.1016/j.apsb.2018.09.007.

Long, J.Z., Svensson, K.J., Bateman, L.A., Lin, H., Kamenecka, T., Lokurkar, I.A., Lou, J., Rao, R.R., Chang, M.R., Jedrychowski, M.P., et al. (2016). The Secreted Enzyme PM20D1 Regulates Lipidated Amino Acid Uncouplers of Mitochondria. Cell 166, 424–435. 10.1016/j.cell.2016.05.071.

Lopez-Otin, C., Blasco, M.A., Partridge, L., Serrano, M., and Kroemer, G. (2013). The hallmarks of aging. Cell 153, 1194–1217. 10.1016/j.cell.2013.05.039.

McNelis, J.C., Lee, Y.S., Mayoral, R., van der Kant, R., Johnson, A.M., Wollam, J., and Olefsky, J.M. (2015). GPR43 Potentiates beta-Cell Function in Obesity. Diabetes 64, 3203–3217. 10.2337/db14-1938.

Mokdad, A.H., Ford, E.S., Bowman, B.A., Dietz, W.H., Vinicor, F., Bales, V.S., and Marks, J.S. (2003). Prevalence of obesity, diabetes, and obesity-related health risk factors, 2001. JAMA 289, 76–79. 10.1001/jama.289.1.76.

Montgomery, M.K., Osborne, B., Brandon, A.E., O’Reilly, L., Fiveash, C.E., Brown, S.H.J., Wilkins, B.P., Samsudeen, A., Yu, J., Devanapalli, B., et al. (2019). Regulation of mitochondrial metabolism in murine skeletal muscle by the medium-chain fatty acid receptor Gpr84. FASEB J 33, 12264–12276. 10.1096/fj.201900234R.

Oh, D.Y., Talukdar, S., Bae, E.J., Imamura, T., Morinaga, H., Fan, W., Li, P., Lu, W.J., Watkins, S.M., and Olefsky, J.M. (2010). GPR120 is an omega-3 fatty acid receptor mediating potent anti-inflammatory and insulin-sensitizing effects. Cell 142, 687–698. 10.1016/j.cell.2010.07.041.

Page, K.A., Williamson, A., Yu, N., McNay, E.C., Dzuira, J., McCrimmon, R.J., and Sherwin, R.S. (2009). Medium-chain fatty acids improve cognitive function in intensively treated type 1 diabetic patients and support in vitro synaptic transmission during acute hypoglycemia. Diabetes 58, 1237–1244. 10.2337/db08-1557.

Pagliarini, D.J., and Rutter, J. (2013). Hallmarks of a new era in mitochondrial biochemistry. Genes Dev 27, 2615–2627. 10.1101/gad.229724.113.

Papamandjaris, A.A., MacDougall, D.E., and Jones, P.J. (1998). Medium chain fatty acid metabolism and energy expenditure: obesity treatment implications. Life Sci 62, 1203–1215. 10.1016/s0024-3205(97)01143-0.

Paschoal, V.A., Walenta, E., Talukdar, S., Pessentheiner, A.R., Osborn, O., Hah, N., Chi, T.J., Tye, G.L., Armando, A.M., Evans, R.M., et al. (2020). Positive Reinforcing Mechanisms between GPR120 and PPARgamma Modulate Insulin Sensitivity. Cell Metab 31, 1173–1188 e1175. 10.1016/j.cmet.2020.04.020.

Poher, A.L., Altirriba, J., Veyrat-Durebex, C., and Rohner-Jeanrenaud, F. (2015). Brown adipose tissue activity as a target for the treatment of obesity/insulin resistance. Front Physiol 6, 4. 10.3389/fphys.2015.00004.

Porter, C., Herndon, D.N., Chondronikola, M., Chao, T., Annamalai, P., Bhattarai, N., Saraf, M.K., Capek, K.D., Reidy, P.T., Daquinag, A.C., et al. (2016). Human and Mouse Brown Adipose Tissue Mitochondria Have Comparable UCP1 Function. Cell Metab 24, 246–255. 10.1016/j.cmet.2016.07.004.

Recio, C., Lucy, D., Purvis, G.S.D., Iveson, P., Zeboudj, L., Iqbal, A.J., Lin, D., O’Callaghan, C., Davison, L., Griesbach, E., et al. (2018). Activation of the Immune-Metabolic Receptor GPR84 Enhances Inflammation and Phagocytosis in Macrophages. Front Immunol 9, 1419. 10.3389/fimmu.2018.01419.

Reilly, S.M., and Saltiel, A.R. (2017). Adapting to obesity with adipose tissue inflammation. Nat Rev Endocrinol 13, 633–643. 10.1038/nrendo.2017.90.

Rosen, E.D., and Spiegelman, B.M. (2014). What we talk about when we talk about fat. Cell 156, 20–44. 10.1016/j.cell.2013.12.012.

Sastre, J., Pallardo, F.V., and Vina, J. (2003). The role of mitochondrial oxidative stress in aging. Free Radic Biol Med 35, 1–8. 10.1016/s0891-5849(03)00184-9.

Schena, G., and Caplan, M.J. (2019). Everything You Always Wanted to Know about beta3-AR * (* But Were Afraid to Ask). Cells 8. 10.3390/cells8040357.

Schilperoort, M., van Dam, A.D., Hoeke, G., Shabalina, I.G., Okolo, A., Hanyaloglu, A.C., Dib, L.H., Mol, I.M., Caengprasath, N., Chan, Y.W., et al. (2018). The GPR120 agonist TUG-891 promotes metabolic health by stimulating mitochondrial respiration in brown fat. EMBO Mol Med 10. 10.15252/emmm.201708047.

Schonfeld, P., and Wojtczak, L. (2016). Short- and medium-chain fatty acids in energy metabolism: the cellular perspective. J Lipid Res 57, 943–954. 10.1194/jlr.R067629.

Shah, R.J., Choudhry, N., and Leiderman, Y.I. (2013). Purtscher-like retinopathy in association with metastatic pancreatic adenocarcinoma and capecitabine therapy. Retin Cases Brief Rep 7, 196–197. 10.1097/ICB.0b013e318280b034.

Shin, H., Ma, Y., Chanturiya, T., Cao, Q., Wang, Y., Kadegowda, A.K.G., Jackson, R., Rumore, D., Xue, B., Shi, H., et al. (2017). Lipolysis in Brown Adipocytes Is Not Essential for Cold-Induced Thermogenesis in Mice. Cell Metab 26, 764–777 e765. 10.1016/j.cmet.2017.09.002.

Simard, J.C., Thibodeau, J.F., Leduc, M., Tremblay, M., Laverdure, A., Sarra-Bournet, F., Gagnon, W., Ouboudinar, J., Gervais, L., Felton, A., et al. (2020). Fatty acid mimetic PBI-4547 restores metabolic homeostasis via GPR84 in mice with non-alcoholic fatty liver disease. Sci Rep 10, 12778. 10.1038/s41598-020-69675-8.

Steinhauser, M.L., Olenchock, B.A., O’Keefe, J., Lun, M., Pierce, K.A., Lee, H., Pantano, L., Klibanski, A., Shulman, G.I., Clish, C.B., and Fazeli, P.K. (2018). The circulating metabolome of human starvation. JCI Insight 3. 10.1172/jci.insight.121434.

Suzuki, M., Takaishi, S., Nagasaki, M., Onozawa, Y., Iino, I., Maeda, H., Komai, T., and Oda, T. (2013). Medium-chain fatty acid-sensing receptor, GPR84, is a proinflammatory receptor. J Biol Chem 288, 10684–10691. 10.1074/jbc.M112.420042.

van Marken Lichtenbelt, W.D., Vanhommerig, J.W., Smulders, N.M., Drossaerts, J.M., Kemerink, G.J., Bouvy, N.D., Schrauwen, P., and Teule, G.J. (2009). Cold-activated brown adipose tissue in healthy men. N Engl J Med 360, 1500–1508. 10.1056/NEJMoa0808718.

Wang, J., Wu, X., Simonavicius, N., Tian, H., and Ling, L. (2006). Medium-chain fatty acids as ligands for orphan G protein-coupled receptor GPR84. J Biol Chem 281, 34457–34464. 10.1074/jbc.M608019200.

Wang, Z., Ning, T., Song, A., Rutter, J., Wang, Q.A., and Jiang, L. (2020). Chronic cold exposure enhances glucose oxidation in brown adipose tissue. EMBO Rep 21, e50085. 10.15252/embr.202050085.

Wellhauser, L., and Belsham, D.D. (2014). Activation of the omega-3 fatty acid receptor GPR120 mediates anti-inflammatory actions in immortalized hypothalamic neurons. J Neuroinflammation 11, 60. 10.1186/1742-2094-11-60.

Williams-Bey, Y., Boularan, C., Vural, A., Huang, N.N., Hwang, I.Y., Shan-Shi, C., and Kehrl, J.H. (2014). Omega-3 free fatty acids suppress macrophage inflammasome activation by inhibiting NF-kappaB activation and enhancing autophagy. PLoS One 9, e97957. 10.1371/journal.pone.0097957.

Wu, J., Bostrom, P., Sparks, L.M., Ye, L., Choi, J.H., Giang, A.H., Khandekar, M., Virtanen, K.A., Nuutila, P., Schaart, G., et al. (2012). Beige adipocytes are a distinct type of thermogenic fat cell in mouse and human. Cell 150, 366–376. 10.1016/j.cell.2012.05.016.

Zoico, E., Rubele, S., De Caro, A., Nori, N., Mazzali, G., Fantin, F., Rossi, A., and Zamboni, M. (2019). Brown and Beige Adipose Tissue and Aging. Front Endocrinol (Lausanne) 10, 368. 10.3389/fendo.2019.00368.

