## Supplemental material for "GPR84-Mediated Signal Transduction Promotes Brown Adipocyte Function"

#### SUPPLEMENTAL INFORMATION

##### Supplemental Figure Legends

**Figure S1. Insensitivity of GPR84 KO mice to HFD-induced obesity and GPR84 distribution in different tissues. Related to Figure 1.**
**(A)** mRNA expression pattern of GPR84 shown in different tissues or organs. n=6-7/group. mRNA levels of GPR84 in different WAT tissues, n=7/group. **(B)** Expression of GPR84 correlated with the expression of UCP1. Data from the GTEx database. **(C)** Glucose levels of GPR84 KO and WT mice on chow at different time points, as determined by GTT (left) and ITT (right) and body weight of the mice. n=12/group. **(D)** Glucose levels of GPR84 KO and WT mice fed a 14-week HFD at different time points, as determined by GTT (left) and ITT (right) and body weight of the mice. n=12/group. One-way ANOVA followed by a Tukey's multiple comparison test. Data are represented as mean±SEM. *P* values determined by two-way ANOVA followed by a Tukey's multiple comparison test (C–D). Data are represented as mean±SEM.

**Figure S2. Body weight, body mass and mitochondrial function in tissues from WT and KO mice at different ages. Related to Figure 2.**
**(A)** Body weight of WT and KO mice. n=10/group. **(B–C)** Body composition analysis including fat mass, lean mass for WT and GPR84 KO mice at different ages. Fat mass and lean mass were normalized to body weight, n=9-10 for old mice (B), n=5 for young mice (C). **(D–G)** $O_2$ consumption rate of gWAT (D) and iWAT (E) from young WT and GPR84 KO mice and gWAT (F) and iWAT (G) of old mice, n=4. **(H–I)** (H) Body weight and (I) tissue weight for eWAT, BAT and liver of WT and KO mice on cold exposure. * $P < 0.05$, ** $P < 0.01$, **** $P<0.0001$. *P* values determined by two-tailed Student's t test for (B–C); two-way ANOVA followed by a Tukey's multiple comparison test (D–G). Data are represented as mean±SEM.

**Figure S3. Gene expressions and OCR in tissues of mice lacking GPR84 on cold exposure. Related to Figure 3.**
**(A)** White selective gene expressions in BAT of WT and KO mice exposed to RT and cold for 6 days, n=8-10/group. **(B–D)** Oxygen consumption rate (OCR) in at basal level and post oligomycin (Oligo), FCCP and rotenone/antimycin-A injection in gWAT (B), iWAT (C) and muscle (D) of mice exposed to cold for 6 days. n = 5 tissues/each time point. **(E–F)** The expressions of mitochondrial-function-related genes (E), and respiratory chain genes (F), n=7-8/group. **(G)** Representative images of Tim23 immunofluorescence of brown adipocytes from WT and KO mice exposed to RT and cold for 6 days, n=3/group. Scale bar indicates 50 µm. **(H)** Serum Triglycerides (left) and NEFA (right) levels of WT and KO mice exposed to RT and cold for 6 days, n=5/group. *P* values determined by two-tailed Student's t test for (A, E, F); two-way ANOVA followed by a Tukey's multiple comparison test (B, C, D, E). Data are represented as mean±SEM.

**Figure S4. Effect of downstream signaling pathway by GPR84 activation on intracellular calcium. Related to Figure 4.**
**(A)** Representative images of UCP1 immunofluorescence staining in brown adipocytes from WT and KO mice, n=3/group. **(B)** Percent of inhibition of CRE-luc in HEK 293 cells transfected with

GPR84 plasmid and CRE-luc plasmid. Cells were incubated with 6-OAU, Embelin, Decanoic acid (C10:0) and Lauric acid (C12:0) for 6h. n=4/each concentration. **(C)** Percent of inhibition of CRE-luc in HEK 293 cells transfected with GPR84 plasmid and SRE-luc. Cells were pre-treated with PTX overnight and incubated with 6-OAU for 6 h. n=4/each concentration. **(D)** Luciferase activity for SRE-Luc in HEK 293 cells transfected with GPR84 SRE-luc and stimulated with 6-OAU, Embelin, Decanoic acid and Lauric acid for 6 h. n=4/each concentration. **(E)** Calcium mobilization for differentiated brown adipocytes treated with or without 6-OAU. After incubation with Fluo-4-AM for 1 h RT, WT brown adipocytes with or without the presence of BAPTA-AM (25 μM). Scale bar indicates 50 μm. Relative fluorescence units were analyzed by ROI of Image J. * $P < 0.05$, ** $P < 0.01$. Two-way ANOVA followed by a Tukey's multiple comparison test (B, C, D). Data are represented as mean±SEM.

**Table S1. Primer sequences for qPCR**

| Gene | Primer forward (5' → 3') | Primer reverse (5' → 3') |
|---|---|---|
| *GPR84* | TCCAATTCTGTCTCCATCCT | CTGACTGGCTCAGATGAAA |
| *UCP1* | AAGCTGTGCGATGTCCATGT | AAGCCACAAACCCTTTGAAAA |
| *Dio2* | TTGGGGTAGGGAATGTTGGC | TCCGTTTCCTCTTTCCGGTG |
| *Cidea* | TCCTATGCTGCACAGATGACG | TGCTCTTCTGTATCGCCCAGT |
| *PGC1α* | TCCTCACACCAAACCCACAGAA | TTGGCTTGAGCATGTTGCGA |
| *PPARγ* | GCCCTTTGGTGACTTTATGGA | GCAGCAGGTTGTCTTGGATG |
| *Cox8b* | TGCTGGAACCATGAAGCCAAC | AGCCAGCCAAAACTCCCACTT |
| *Prdm16* | ACACGCCAGTTCTCCAACCTGT | TGCTTGTTGAGGGAGGAGGTA |
| *Resistin* | AAGAACCTTTCATTTCCCCTCCT | GTCCAGCAATTTAAGCCAATGTT |
| *Leptin* | ATGTGCCCTTCCGATATACAACC | CGTGTCATCCACTAATCTTCTGG |
| *Tie3* | GAGACTGAACACAATCCTAGCC | GGAGTCCACGTACCCCGAT |
| *GSTA3* | AGATCGACGGGATGAAACTGG | CAGATCCGCCACTCCTTCT |
| *Zfp423* | CAGGCCCACAAGAAGAACAAG | GTATCCTCGCAGTAGTCGCACA |
| *MCP-1* | TTAAAAACCTGGATCGGAACCAA | GCATTAGCTTCAGATTTACGGGT |
| *Il6* | CCAGAGATACAAAGAAATGATGG | ACTCCAGAAGACCAGAGGAAAT |
| *TNFα* | GCCACCACGCTCTTCTGCCT | GGCTGATGGTGTGGGTGAGG |
| *IL-1β* | AAATACCTGTGCCCTTGGGC | CTTGGCATCCACACTCTCCAG |
| *Cycs* | CCAGGTATACAAGCAGGTGTGCTC | CATCATTAGGGCCATCCTGGAC |
| *Cox2* | AGTTGATAACCGAGTCGTTCTGCCA | TCGGCCTGGGATGGCATCAGT |
| *Cox4* | ATTGGCAAGAGAGCCATTTCTAC | CACGCCGATCAGCGTAAGT |
| *Cox6c* | GGGAAGGACGTTGGTGTAGA | CTTATAGGCAGCGGCAACTC |
| *Cox8a* | TTCCTGCTTCGTGTGTTGTC | GATTGCAGAAGAGGTGACTGG |
| *Nd1* | CAGCCGGCCCATTCGCGTTA | AGCGGAAGCGTGGATAGGATGC |
| *Nd2* | TCCTCCTGGCCATCGTACTCAACT | AGAAGTGGAATGGGGCGAGGC |
| *Nd3* | ACCCTACAAGCTCTGCACGCC | GCTCATGGTAGTGGAAGTAGAAGGGCA |
| *Nd4* | TCGCCTACTCCTCAGTTAGCCACA | TGATGATGTGAGGCCATGTGCGA |
| *Nd5* | TCGGAAGCCTCGCCCTCACA | AGTAGGGCTCAGGCGTTGGTGT |
| *Nd6* | AATACCCGCAAACAAAGATCACCCAG | TGTTGGGGTTATGTTAGAGGGAGGGA |
| *Atp6* | GCTCACTCGCCCACTTCCTTCC | GCCGGACTGCTAATGCCATTGGTT |
| *Atp8* | ATGCCACAACTAGATACATCAACA | GGGGTAATGAATGAGGCAAA |
| *RPL19* | ATGAGTATGCTCAGGCTACAGA | GCATTGGCGATTTCATTGGTC |

**Table S2. Values of Log [IC50] and Log [EC50] for luciferase assay. Related to Figure S4.**

| Figures | Log [IC50] | Log [EC50] |
|---|---|---|
| Figure S2B | 6-OAU: -7.868±0.0569<br>Embelin: -7.577±0.466<br>Lauric acid:-6.669±0.168<br>Decanoic acid:-7.122±0.360 | N/A |
| Figure S2C | Empty vendor+6-OAU: -6.301±0.2125<br>GPR84+6-OAU: -9.078±0.8645<br>GPR84+6-OAU+PTX:-7.474±0.1204 | N/A |
| Figure S2D | N/A | 6-OAU: -6.842±0.8907<br>Embelin: -0.4908±10166<br>Lauric acid:-65.559±0.7406<br>Decanoic acid:-7.925±0.2949 |

### Figure S1. Sun et al. — Related to Figure 1

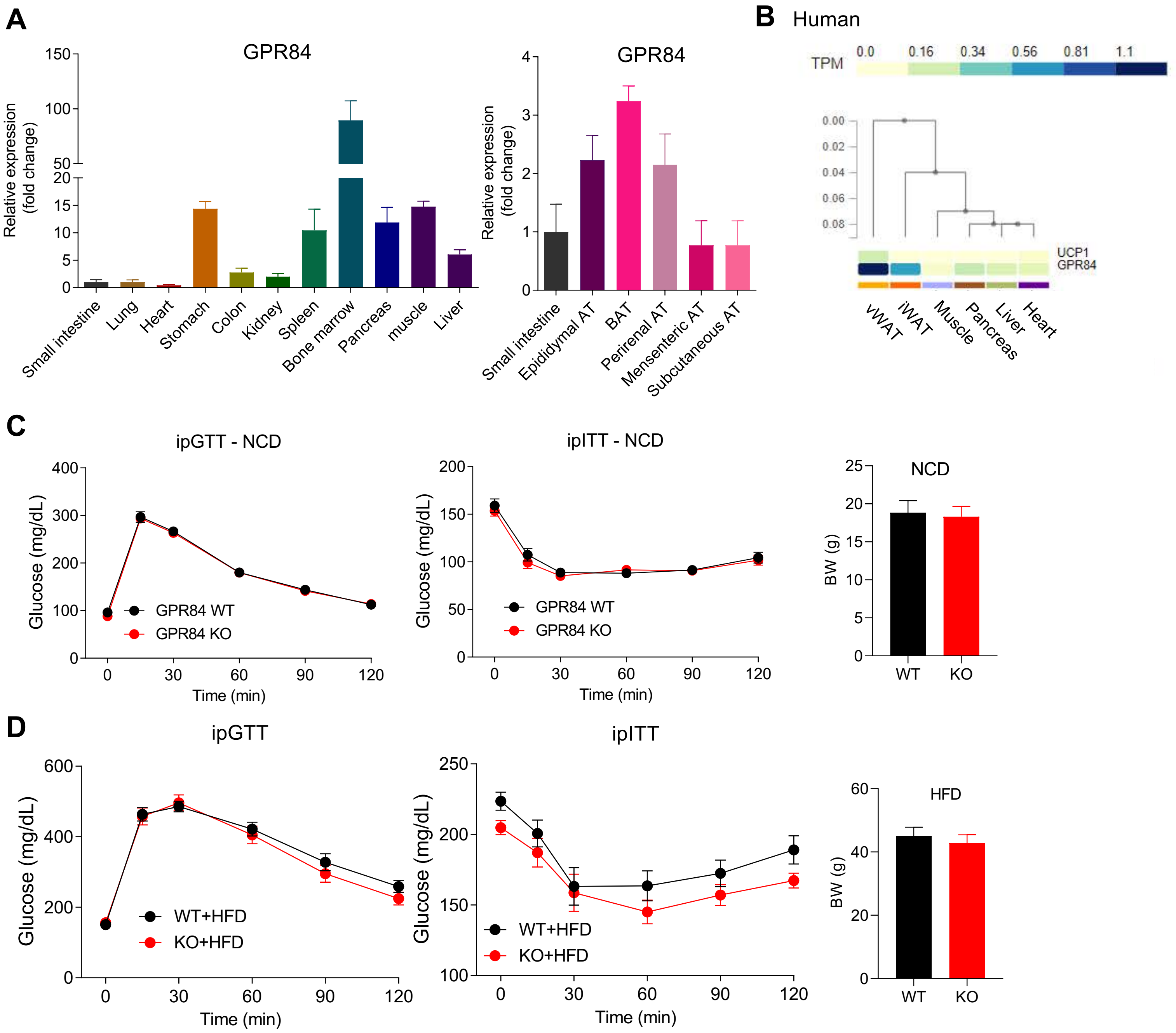

### Figure S2. Sun et al. — Related to Figure 2

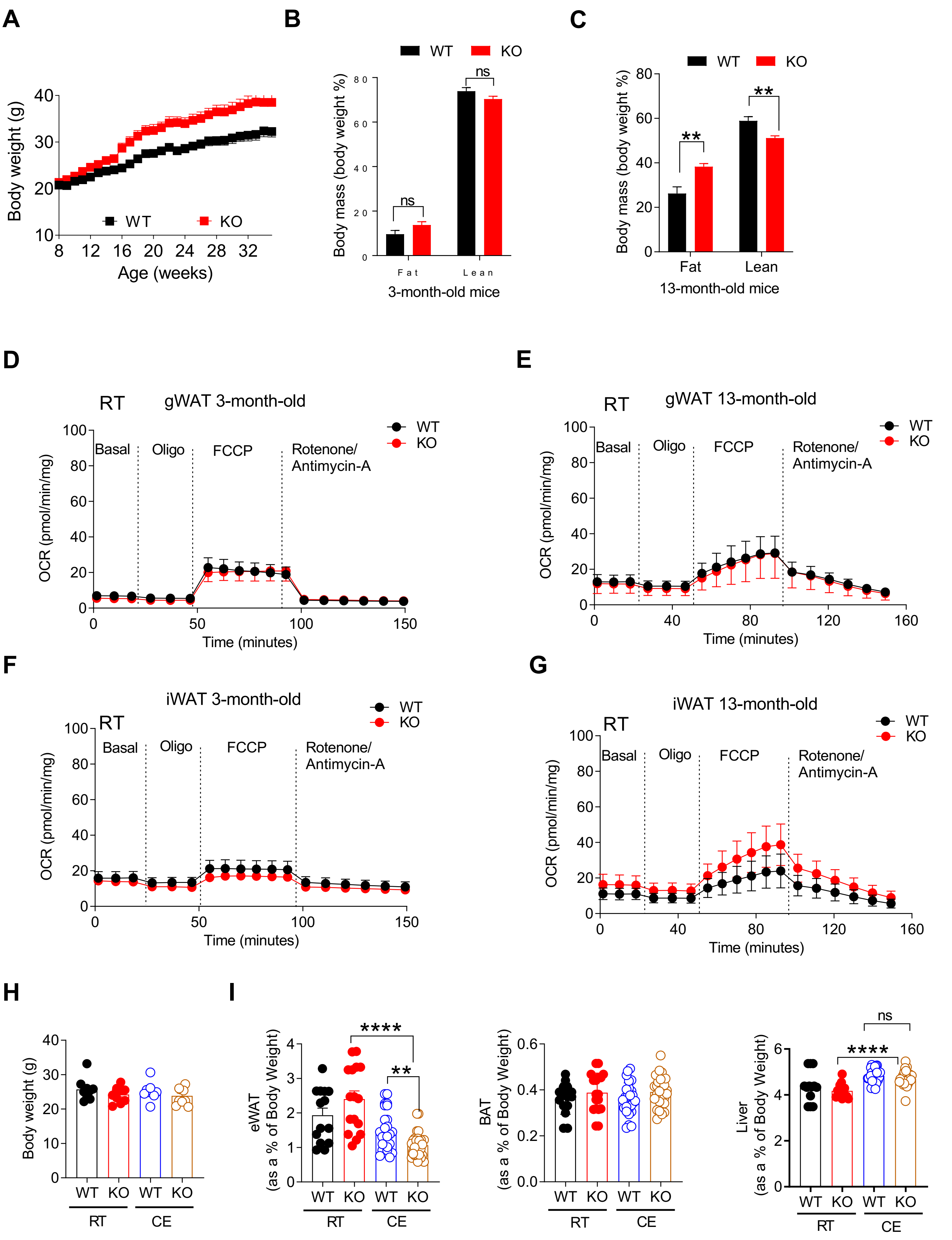

### Figure S3. Sun et al. — Related to Figure 3

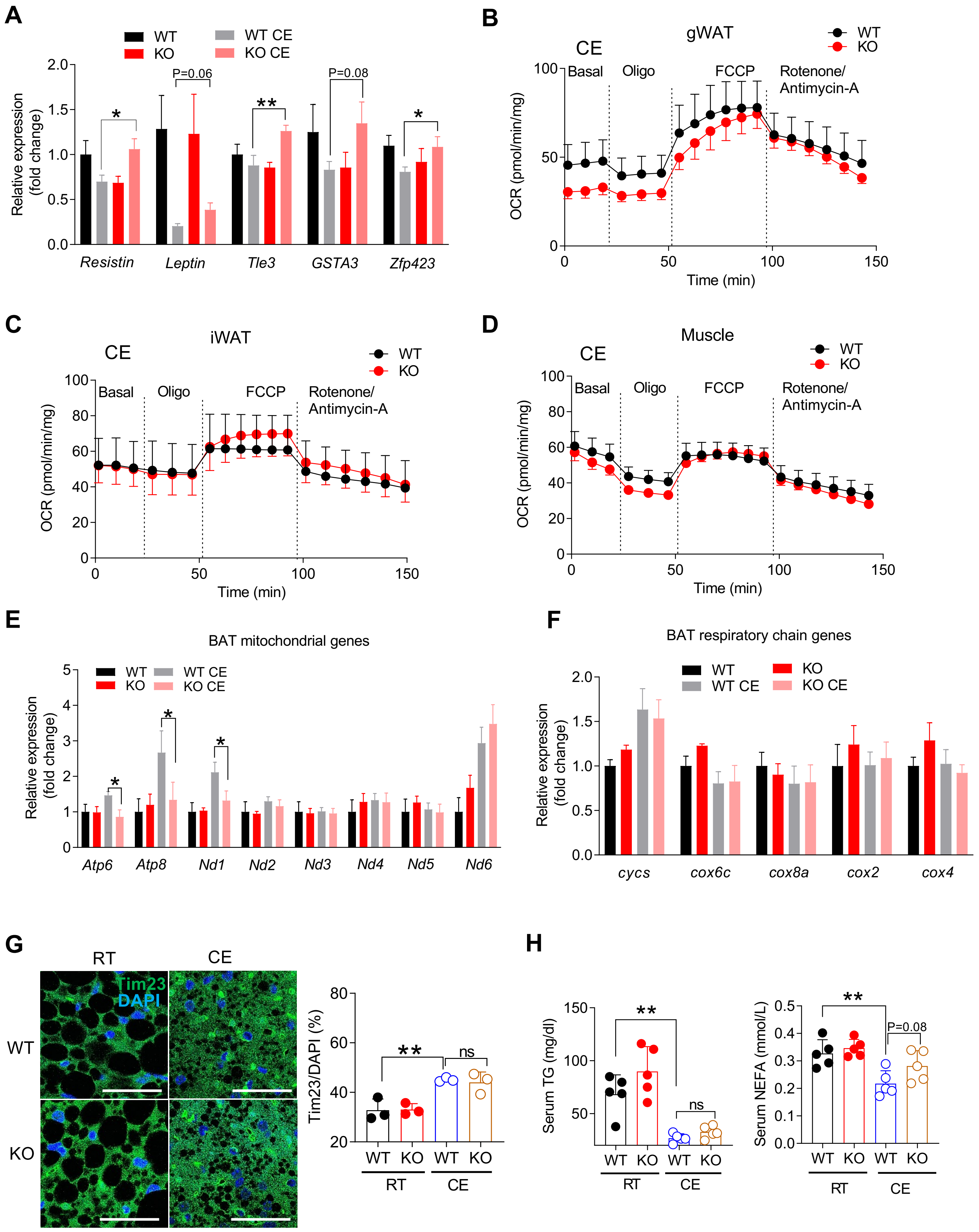

**Figure S4. Sun et al. — Related to Figure 3 and Table S2**

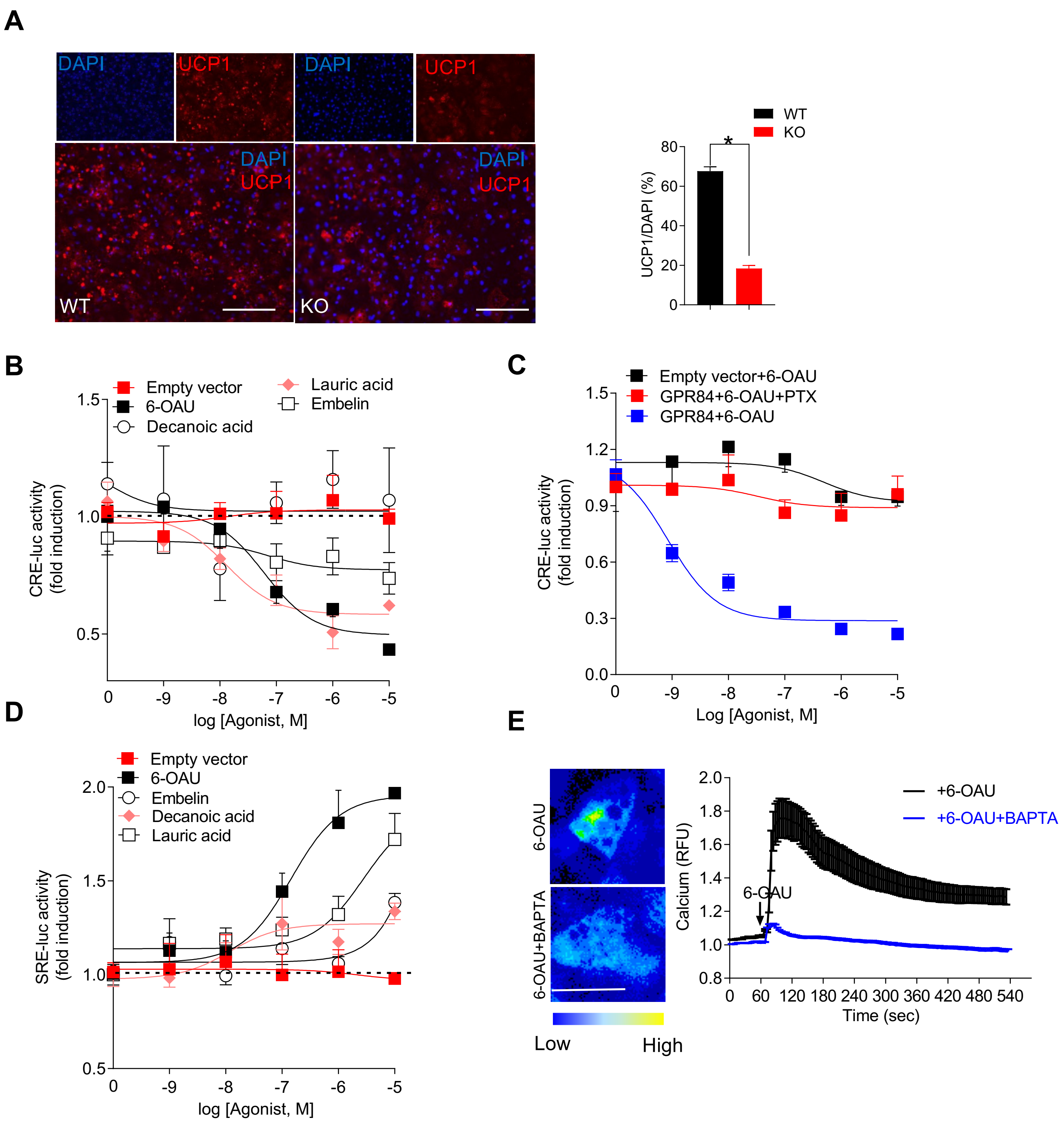
